# Social selectivity and social motivation in voles

**DOI:** 10.1101/2021.07.30.454556

**Authors:** Annaliese K. Beery, Sarah A. Lopez, Katrina L. Blandino, Nicole S. Lee, Natalie S. Bourdon, Todd H. Ahern

## Abstract

Selective relationships are fundamental to humans and many other animals, but relationships between mates, family members, or peers may be mediated differently. We examined connections between social reward and social selectivity, aggression, and oxytocin receptor signaling pathways in rodents that naturally form enduring, selective relationships with mates and peers (prairie voles) or peers (meadow voles). Female prairie and meadow voles worked harder to access familiar vs. unfamiliar individuals, regardless of sex, and huddled extensively with familiar subjects. Male prairie voles also displayed strongly selective huddling preferences for familiar animals, but worked hardest to repeatedly access females vs. males, with no difference in effort by familiarity. This demonstrates a fundamental disconnect between motivation and social selectivity in males, and reveals a striking sex difference in pathways underlying social monogamy. Meadow voles exhibited social preferences but low social motivation, consistent with tolerance rather than reward supporting social groups in this species. Natural variation in oxytocin receptor genotype was associated with oxytocin receptor density, and both genotype and receptor binding predicted individual variation in prosocial and aggressive behaviors. These results provide a basis for understanding species, sex, and individual differences in the mechanisms underlying the role of social reward in social preference.

## INTRODUCTION

The brain regions and neurochemicals involved in social behaviors show remarkable conservation across species (O’Connell and Hofmann, 2011). At the same time, social behavior is not a unified construct, with different species exhibiting distinct social structures and behavioral repertoires. The formation of selective social relationships is a particular hallmark of both human and prairie vole societies. Such relationships are difficult to study in traditional lab rodents because mice, rats, and other rodents typically don’t form preferences for known peers or mates (Triana-Del Rio et al., 2015; Schweinfurth et al., 2017; Beery et al., 2018; Cymerblit-Sabba et al., 2020; Insel et al., 2020; Beery and Shambaugh, 2021). In species that form specific relationships, selectivity may be based on reward and prosocial motivation towards specific individuals or on avoidance (fear, aggression) of unfamiliar individuals. The role of social motivation and tolerance may also differ by familiarity, sex, and type of relationship (e.g., same-sex peer versus opposite-sex mate). Voles provide an opportunity to probe the role of selectivity and social reward across relationship types and social organization.

The reinforcing properties of social interaction have been demonstrated in a variety of rodent species and contexts, often through conditioned place preference for a socially associated environmental cue (e.g. Panksepp and Lahvis, 2007; Dölen et al., 2013; Goodwin et al., 2019). Operant conditioning for access to a social stimulus has been used to more directly measure motivation for specific types of social interaction, particularly access to pups, social play, and sexual opportunities (reviewed in Trezza et al., 2011). Social motivation has also been assessed with access to novel same-sex peers (Martin and Iceberg, 2015; Achterberg et al., 2016; Borland et al., 2017). Often social interactions are affiliative, but in some contexts animals will work for access to aggressive interactions (Azrin et al., 1965; Falkner et al., 2016; Golden et al., 2017). To date, only one study has examined the role of familiarity in social motivation, in novelty-preferring female rats (Hackenberg et al., 2021), and none have done so with mate relationships.

Prairie voles, *Microtus ochrogaster,* and meadow voles, *Microtus pennsylvanicus,* both form selective social relationships but exhibit different social organization and mating systems. Prairie voles are socially monogamous, forming long-term selective relationships between males and females that have been studied for decades (Carter et al., 1995; Walum and Young, 2018). Prairie voles also form selective relationships with familiar same-sex cage-mate “peers” (DeVries et al., 1997; Beery et al., 2018; Lee et al., 2019). Meadow voles are promiscuous breeders that transition to living in social groups and sharing nests during winter (Getz, 1972; Madison and McShea, 1987). Under conditions of short day length in the laboratory, female meadow voles exhibit greater social huddling and less aggression than their long day length counterparts (Beery et al., 2008b; Lee et al., 2019). These vole species thus allow comparison of the properties of peer relationships across species (prairie vole peers vs. meadow vole peers) and relationship type within species (prairie vole mates vs. prairie vole peers).

Prairie voles exhibit socially conditioned place preferences (sCPP) for familiar opposite-sex peers (Ulloa et al., 2018; Goodwin et al., 2019), and in some circumstances for same-sex peers (Lee and Beery, 2021). In contrast, meadow voles do not form sCPP, and may even condition away from social cues (Goodwin et al., 2019). Neurochemical pathways underlying social reward also vary between species and relationship type; dopamine is necessary for the formation of opposite-sex pair bonds in prairie voles (Aragona and Wang, 2009), but is not necessary for the formation of same-sex peer preferences in meadow or prairie voles (Beery and Zucker, 2010; Lee and Beery, 2021). These initial findings suggest that social selectivity may result from differential social motivation and tolerance in these species.

Voles demonstrate striking preferences for familiar vs. novel peers and mates, assessed using the partner preference test (Williams et al., 1992b; Beery, 2021). This test quantifies preference, but as no effort is required to access a conspecific, it cannot distinguish between prosocial motivation and avoidance of unfamiliar conspecifics. To examine the role of motivation in relationships, we assessed effort expended by voles of different sexes (male, female), relationship types (same-sex, opposite-sex), and species (prairie vole, meadow vole) to reach social targets in an operant conditioning paradigm. Subjects underwent >60 active training and testing days (Figure 1). Responses (lever presses) in lightly food-restricted voles were shaped and reinforced using a food reward, followed by 8 days of pressing for a food reward on a progressive ratio 1 (PR-1) schedule. Social testing consisted of 8 days of testing in which rewards yielded 1 minute of access to the familiar (same or opposite-sex) partner, and 8 days with access to a sex-matched stranger (order balanced within groups). We assessed effort expended to access familiar and novel social stimuli in four groups of prairie voles (Figure 1): females lever pressing for a female conspecific (F➤F), females pressing for a male conspecific (F➤M), males pressing for a male conspecific (M➤M), and males pressing for a female conspecific (M➤F). Meadow vole females (F➤F) were also trained and tested for 8 days of familiar and 8 days of novel vole exposure, counterbalanced. A subset of voles was used to explore the reward value of an empty chamber, extinguishing timelines, and relationships between oxytocin receptor density, oxytocin receptor genotype, and behavior.

**Figure 1.**
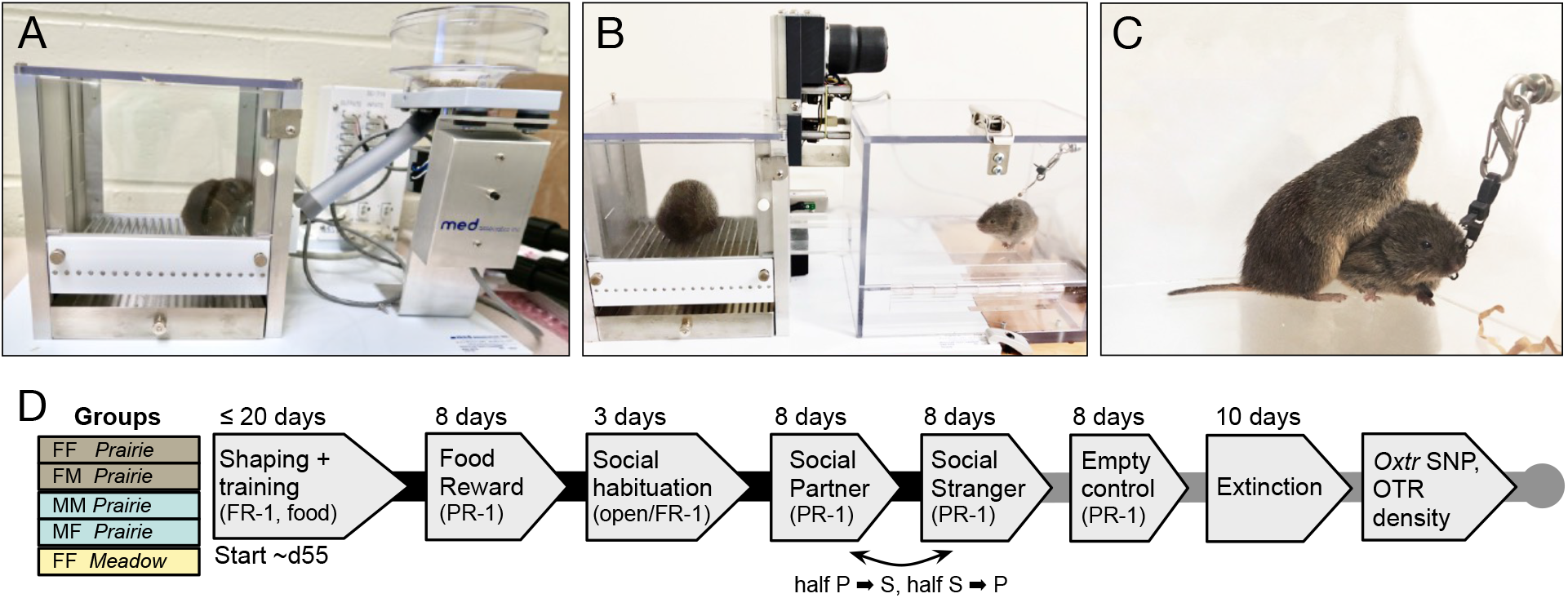
Overview of apparatuses, timeline, and testing groups. **A.** Lever pressing in voles was shaped and trained using food reinforcement. **B,C.** In social operant testing a lever operated a motorized door, providing 1 min access to a conspecific tethered in a connected compartment. **D.** Five groups were tested, abbreviated here as focal sex-partner sex-species abbreviation (e.g., FF *Prairie* indicates a female prairie vole trained as a lever presser and housed with a female partner). Prairie = prairie vole (*Microtus ochrogaster*); Meadow = meadow vole (*Microtus pennsylvanicus*). Black lines connect testing phases completed by all study subjects; gray lines connect additional phases completed by a subset of subjects.

Oxytocin is involved in social recognition as well as in preference for familiar individuals (reviewed in Anacker and Beery, 2013), and in many instances, oxytocin signaling alters the rewarding properties of social stimuli (Dölen et al., 2013; Borland et al., 2019). Genotype at the intronic NT213739 single nucleotide polymorphism (SNP) in the oxytocin receptor gene (<u>*Oxtr*</u>) has recently been associated with individual variation in striatal oxytocin receptor density as well as partner preference formation in prairie voles (King et al., 2016; Ahern et al., 2021). We assessed *Oxtr* genotype at this SNP for all prairie voles at the conclusion of testing and conducted receptor autoradiography to assess variation in neural oxytocin receptor density in female prairie voles.

Together these studies allowed us to examine how the reward value of social contact differs between male and female prairie voles, between opposite-sex and same-sex pairings, and between meadow and prairie vole FF pairings. We found both similarities in and striking differences between social motivation across species, sexes, and pairing types. Detailed examination of social behaviors during social access further underscored the distinction between social motivation and familiarity preference, especially in males. In addition to these group differences in social motivation, individual differences in oxytocin receptor genotype predicted variation in oxytocin receptor density, and both genotype and receptor density related to aggressive and prosocial behaviors.

## RESULTS

### Sex-specific patterns of social effort in prairie voles

Males and females showed qualitatively different response patterns, so responses in the social chambers were analyzed separately (Beltz et al., 2019). Two-way RM-ANOVA was performed with familiarity of the tethered stimulus (partner/stranger) as the within-subjects/repeated measure, and sex of the tethered stimulus (opposite-sex/same-sex) as a between-subjects measure. Female prairie voles pressed more for familiar partners than unfamiliar strangers, with no effect of opposite-sex vs. same-sex pairings (Figure 2A, effect of stimulus familiarity: F_(1, 14)_ = 15.17, *p =* 0.0016, effect of stimulus sex: F_(1, 14)_ = 0.44, *p =* 0.51, subject matching: F_(14, 14)_ = 4.2, *p =* 0.0057, no significant interaction). Paired *t*-tests were used for within-group comparisons of responses for the partner or stranger: familiarity preferences were significant in females paired with males (t_(7)_ = −2.7, *p =* 0.03) as well as in females paired with females (t_(7)_ = - 4.1, *p =* 0.0048).

**Figure 2.**
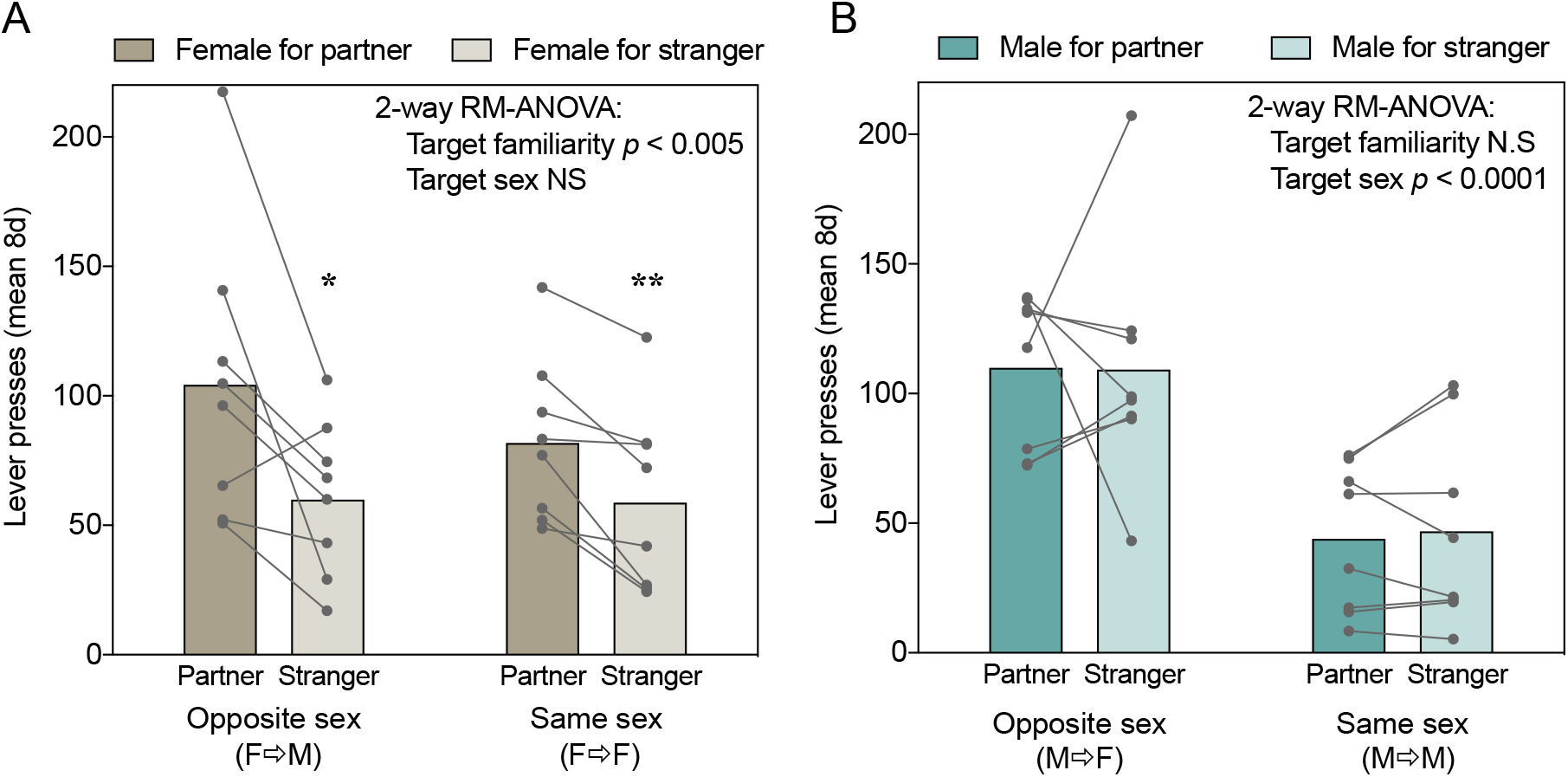
Sex-specific patterns of effort expended to access different social stimuli. **A.** Female prairie voles responded more for familiar than unfamiliar voles of either sex. **B.** Male prairie voles pressing for females responded more than did males pressing for males, regardless of familiarity. Dots represent mean number of responses across eight 30-minute PR-1 sessions for each vole. Bars represent group means. Asterisks indicate significant familiarity preferences within groups (paired *t*-tests). * = *p*<0.05, ** = *p*<0.01.

Male prairie voles pressed at a higher rate for opposite-sex social stimuli regardless of familiarity (effect of stimulus sex: F_(1, 14)_ = 17.4, *p =* 0.0009; effect of familiarity, F_(1, 14)_ = 0.013, *p =* 0.91, no significant effects of subject matching or interaction).

Because each vole was tested in 8 consecutive sessions of each type, familiarity preference could also be assessed within individuals across days. Significant within-vole familiarity preferences were present in more female pressers (6/8 F➤M and 3/8 F➤F) than males (1/8 M➤F and 0/8 M➤M pairs) (supplemental data Figure 1; *p =* 0.0059 fisher’s exact test). One male in a M➤F pair exhibited a significant preference for stranger females (Figure S1), and mounted/copulated with strangers in multiple test sessions.

### Social motivation and behavior were parallel in female but not male prairie voles

In female prairie voles, the familiarity preference for both mates and peers in lever pressing was mirrored in cohabitation time and huddling. Even when these behaviors were scaled relative to lever presses (and thus access time), females spent a significantly higher fraction of the available time in the social chamber (time in social chamber/access time) when it was occupied by a familiar vole rather than a novel one (effect of familiarity F_(1,14)_=95.06, p<0.0001; subject matching F_(14,14)_ = 2.789, *p =* 0.03; others NS; two-way RM-ANOVA, Figure 3A). Females also spent more of the available time huddling (time spent in immobile side-by-side contact/access time) with familiar rather than unfamiliar conspecifics of either sex (effect of familiarity: F_(1,14)_ = 25.82, *p =* 0.0002; others N.S.; two-way RM-ANOVA, Figure 3C). Within-group matched comparisons of time spent with a partner or stranger also revealed that females exhibited significant familiarity preferences in time spent in the social chamber or huddling with the stimulus animal relative to time with access (time in social chamber/access time: FF: *p <* 0.0001, FM: *p =* 0.0006; time huddling/access time: FF *p =* 0.0090; FM *p =* 0.0083; paired *t*-tests).

**Figure 3.**
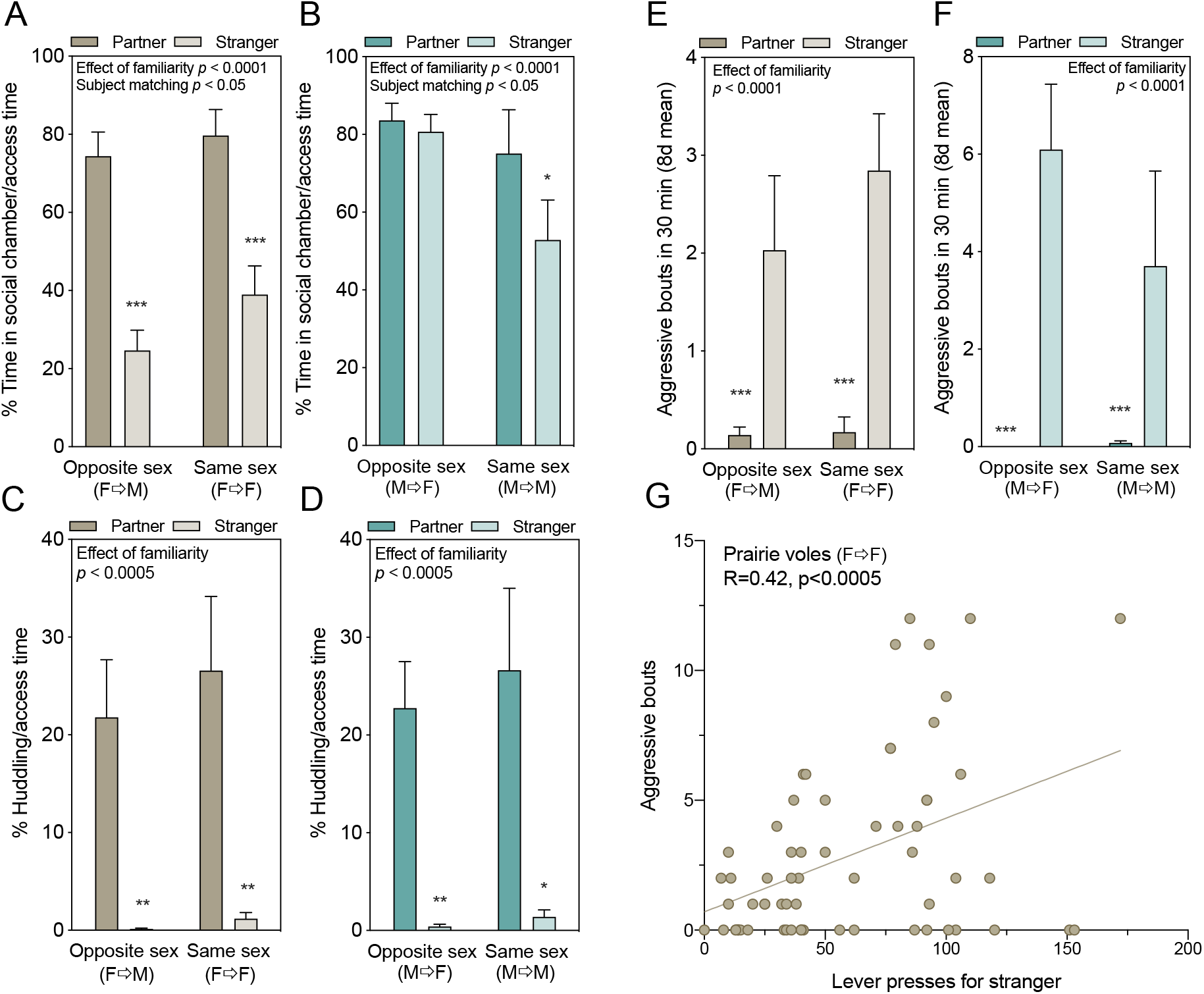
Affiliative and aggressive interactions with stimulus voles. Data represent the 8-day testing mean for each vole (n = 8/group, ± SEM). **A,B.** Percent of time focal voles spent in the social chamber relative to time when the door was open, allowing chamber access. Females shown in A, males in B. **C,D.** Percent time spent huddling out of access time (i.e. when the door was raised). Significant effects of two-way RM-ANOVA are reported above each graph. Asterisks represent the results of within-groups paired *t*-tests. **E,F**. Prairie voles exhibited significantly more bouts of aggression towards strangers (*p <* 0.0001), and there were no significant effects of sex of the presser or of the social target. **G**. Female prairie vole lever pressing for access to an unfamiliar stranger was associated with bouts of aggression toward that stranger (R = 0.42; *p <* 0.0005), although this effect disappears if scaled by access time. No relationship between stranger lever pressing and aggression was present in female prairie voles tested with males, or males tested with either sex. * *p <* 0.05, ** *p <* 0.01, *** *p <* 0.001.

In contrast, while males exhibited no familiarity preferences in lever pressing responses, they exhibited strong familiarity preferences in social interaction. Males spent more of the available time in the social chamber when the tethered stimulus was familiar (effect of familiarity F_(1,14)_ = 6.33, *p =* 0.02; subject matching F_(14,14)_ = 4.459, *p =* 0.0042; others NS; 2-way RM-ANOVA, Figure 3B), and huddling behavior was even more specific, with a strong effect of stimulus familiarity (partner vs. stranger) and no effect of stimulus sex (opposite- vs. same-sex) on the percent of [time huddling]/[time with access to the social chamber] (effect of familiarity: F_(1,14)_ = 25.27, *p =* 0.0002; all else N.S.; Figure 3D). Within-group matched comparisons also revealed significant familiarity preferences in huddling time relative to access (huddling/access time: MM: p = 0.0177, MF *p =* 0.0022), with lesser or no familiarity preference in chamber time (social chamber/time with access: MM: *p =* 0.0390, MF: *p =* 0.56; paired *t*-tests). There was no apparent sex difference in huddling behavior between male and female prairie voles, confirmed by pooling males and females in a 3-way ANOVA (effect of focal sex NS, *p =* 0.91; significant effect of stimulus familiarity (F_(1,56)_ = 48.03, *p <* 0.0001); effect of stimulus sex NS; no significant interactions).

### Other social/sexual behaviors in prairie voles

Aggressive behavior was exhibited by prairie voles in all groups and was analyzed by RM-ANOVA on all voles tested with partners and strangers (between subjects factors: sex of presser (M/F)*pairing type [same/opposite sex]; within subjects factor: target familiarity). Both males and females engaged in far more bouts of aggression with strangers than familiar partners (F_(1,29)_ = 30.22, *p <* 0.0001, Figure 3 E,F). There was no significant effect of sex of the presser (F_(1, 29)_ = 3.36, *p =* 0.077), pairing type (same-sex or opposite-sex), or interactions between these variables.

Because aggression was primarily targeted at strangers, we assessed whether greater lever pressing for strangers was associated with aggression. There were no relationships between stranger-directed lever pressing and aggression in male prairie voles, or in females pressing for males. In female prairie voles pressing for unfamiliar females (F➤F), there was a moderate but significant correlation between stranger-directed pressing and aggressive bouts (R = 0.42, *p =* 0.0005). Unlike effects reported for huddling/access time, however, this effect disappears when scaled by access time, suggesting this phenomenon represents opportunistic aggression towards the stranger when the motorized door is raised.

Mounting behavior was present in five prairie voles, all of which were male prairie voles tested with novel (unfamiliar) female voles. This distribution was significantly non-random across the eight testing combinations used in prairie voles (e.g. male with female partner, male with female stranger, etc.) (χ^2^_(7)_ = 37.97, *p <* 0.0001). These five voles exhibited an average of 6 bouts of mounting per testing session.

### Neural oxytocin receptor density related to behavior, housing, and genotype

Oxytocin receptor density (OTR) was associated with both motivated and aggressive social behaviors in different brain regions in female prairie voles (males not assayed). There was a strong positive correlation between oxytocin receptor density and lever presses for same-sex partners in the nucleus accumbens (NAcc) core (R = 0.959, *p =* 0.0098) and shell (R = 0.948, *p =* 0.0141, Figure 4A). There was also a strong positive correlation between mean bouts of aggression and OTR density in the bed nucleus of the stria terminalis (BNST) in female prairie voles (R = 0.719, *p =* 0.0126), again connecting receptor binding to behavior.

**Figure 4.**
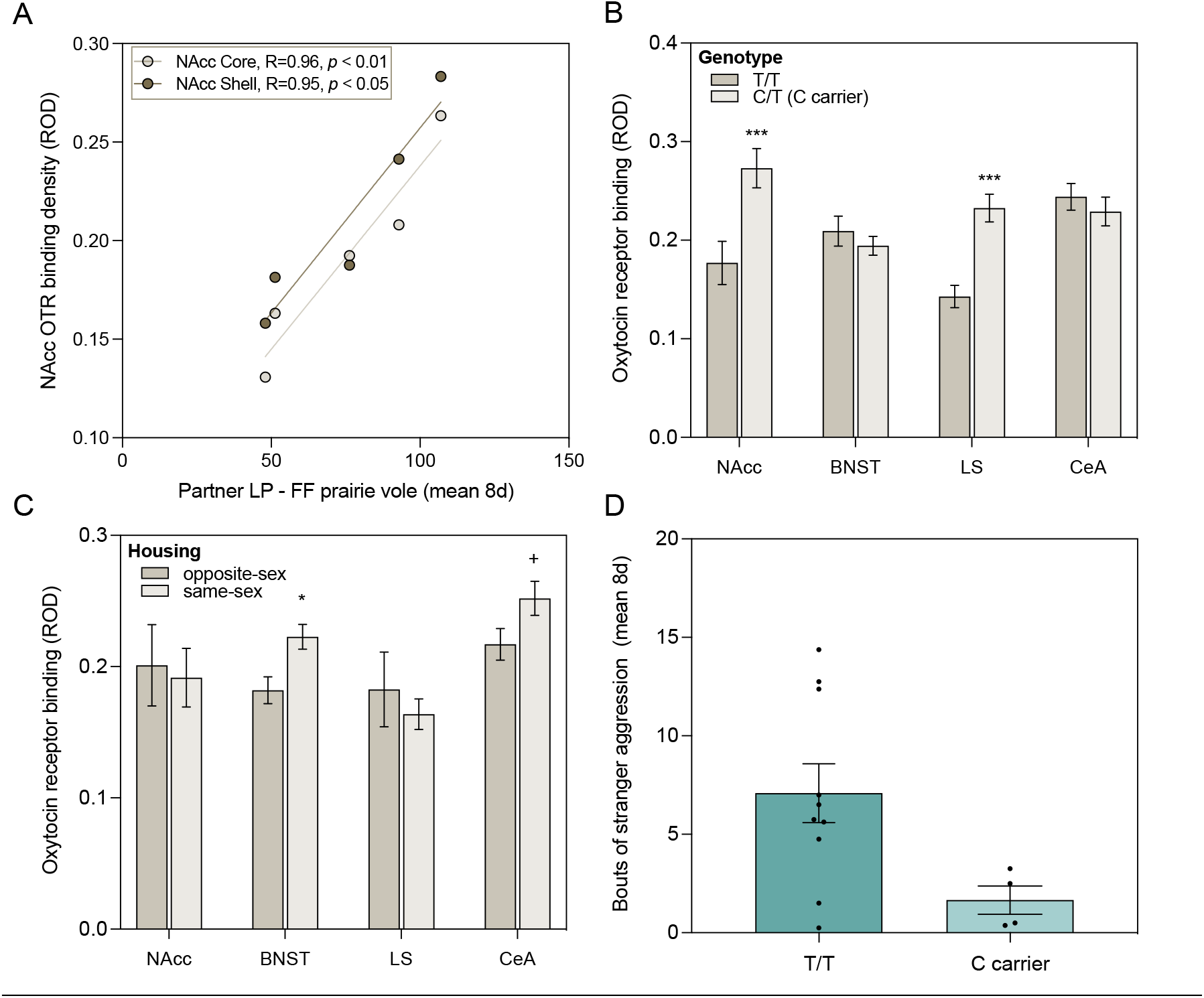
Oxytocin receptor genotype, oxytocin receptor density, and behavior. **A.** OTR binding in the NAcc core and shell were strongly correlated with individual variation in lever pressing for a partner. **B.** Individual variation in OTR density was influenced by genotype in multiple brain regions, with higher OTR expression in the NAcc and LS of C carriers. **C.** OTR density also varied in response to housing/pairing condition (opposite-sex vs. same-sex), in distinct regions from those influenced by genotype. **D.** OTR genotype at the NT213739 SNP significantly predicted behavior across housing conditions, with the greatest effect on stranger-directed aggression in male prairie voles.

Oxytocin receptor density varied with both genotype and housing. The genotyped study population was comprised of C/C (1), C/T (9), and T/T (18) individuals; individuals with one or two C alleles were reported together as ‘C carriers’ as in prior studies (Ahern et al., 2021). C carriers exhibited higher OTR binding in specific brain regions (two-way ANOVA; effect of genotype: *p <* 0.002, effect of brain region: *p <* 0.0234, genotype*region interaction: *p <* 0.0008). In particular, C carriers exhibited higher binding in the NAcc and lateral septum (LS) (*p <* 0.0002 and *p <* 0.0005, adjusted for multiple comparisons; Figure 4B). Housing also influenced OTR density, but in different brain regions. Females housed with same-sex cage-mates showed no difference in OTR density in the NAcc or LS, higher OTR density in the BNST (*t*(8.99) = 2.93, *p* = 0.0167), and a non-significant trend in the central amygdala (CeA) (*t*(8.71) = 1.92, *p =* 0.0883) compared to females housed with opposite-sex cage-mates (Figure 4C).

### Genotype-behavior connections

Genotype was randomly assorted across groups, so comparisons to behavior were made when the samples could be pooled. Stranger-directed aggressive behavior was consistent across pairing type, and was thus compared in all males and all females. Male C carriers exhibited far fewer bouts of aggression than T/T individuals (*p <* 0.0068, Figure 4D), with no effect of genotype found in females (*p =* 0.993). Lever pressing effort varied by pairing type in males, preventing pooling, but females could be pooled; female C carriers exhibited a trend towards greater lever pressing for the partner/total (*p =* 0.056). This is consistent with both higher NAcc OTR in C carriers, and the positive correlation between NAcc OTR and partner lever pressing.

### Interspecific comparisons: responses were reward-specific and comparable across species and sexes

There were no sex or species differences in the number of lever pressing responses for a food reward (PR-1 schedule; 8 days averaged per subject) between female prairie voles, male prairie voles, and female meadow voles (F_(2,40)_ = 1.18, *p =* 0.32; one-way ANOVA; Figure 5A). Food responses and social responses were converted to response rates for comparison across trials with different active lever pressing periods: individual response rates for a food reward did not predict response rates during social testing for either the partner (*p =* 0.78) or the stranger (*p =* 0.98), indicating that responses were not subject-specific across reward types (Figure 5B). These findings validate the specificity of comparisons across species, sexes, and reward types.

**Figure 5.**
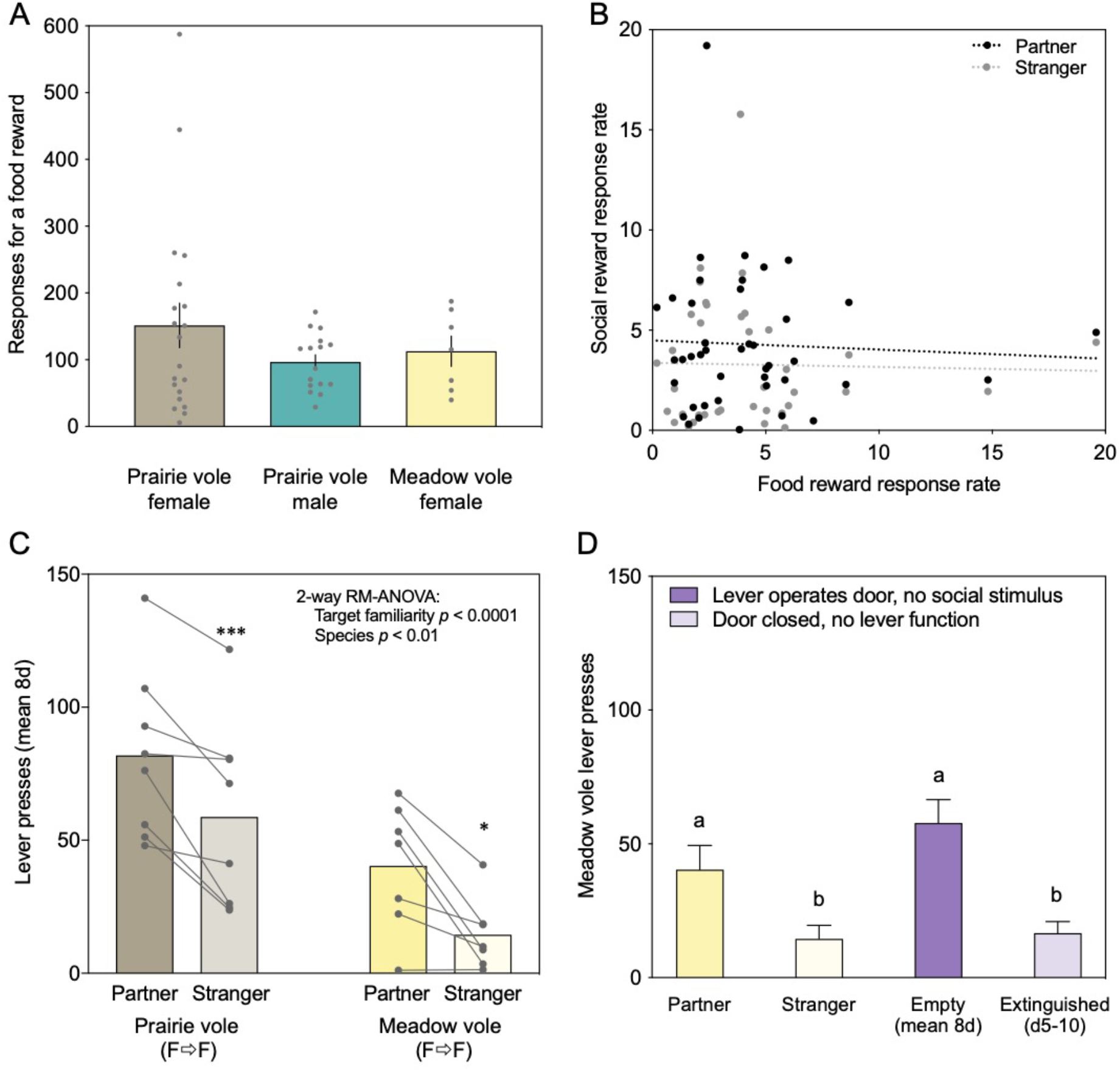
Quantifying responses across species, sexes, and reward types. **A.** Responses did not significantly differ between prairie voles of different sexes or between meadow and prairie vole females. Each data point represents the 8-day mean of responses from a vole tested using a PR-1 schedule in 30-min sessions. **B.** Food response rate did not predict social response rate for familiar or unfamiliar stimuli. Data points show prairie vole response rates for food pellets on a PR-1 schedule (8 day mean for each vole) versus social reward (black: partner; gray: stranger) on a PR-1 schedule (8 day mean for each vole). **C.** Meadow voles, like prairie voles, pressed more for a partner than a stranger, but pressed significantly less overall. **D.** Social pressing for a partner in meadow voles was no higher than pressing for an empty chamber, and stranger pressing was similar to the minimum achieved by extinction.

### Meadow voles exhibited familiarity preferences but low social response rates

Female meadow voles pressed significantly more for familiar females than novel females (t_(6)_=3.637, *p*=0.0109, paired *t*-test, Figure 5C; males not tested). This preference was individually significant within four of the seven meadow voles (Supplementary Figure S2). Comparisons of time spent with a partner or stranger when the door was up also revealed significant familiarity preferences (P vs. S for social chamber/access time: *p =* 0.0351; P vs. S for huddling/access time: *p =* 0.0357; paired *t*-tests).

Despite familiarity preference, meadow vole response rate for both partners and strangers was low. Direct comparison with female prairie voles tested under the same conditions reveals that while both groups pressed more for familiar partners than for strangers, there was significantly less lever pressing in female meadow voles (2-way ANOVA, effect of target familiarity: *p <* 0.001, effect of species: *p <* 0.01, Figure 5C). Comparison of lever presses between social conditions and non-social “empty control” conditions indicates that, for female meadow voles, the partner was not more rewarding than the empty chamber control, stranger pressing was significantly lower than empty control, and it was similar to the post-extinction level of pressing (Figure 5D).

### Other social/sexual behaviors in meadow voles

Aggression was rare in meadow vole trials (mean 0.3 bouts/trial), and as in our prior studies (Lee et al., 2019) it was significantly less frequent than aggression between female prairie voles (mean 2.3 bouts/trial, species difference: *t*(3.83), *p*=0.001). No mounting behavior was observed in meadow vole tests, all of which were conducted in female voles.

### Empty chamber control and extinction

At the conclusion of social testing, a subset of each group was tested for effort expended to explore an empty chamber without a tethered partner or stranger for 8 days each (n=7 meadow females, 20 prairie females: 10 housed FM and 10 housed FF, and 16 prairie males: 8 housed MM and 8 housed MF). There was no species difference in pressing for the empty chamber (meadow vole female vs. prairie vole female). In both male and female prairie voles, the extent of lever pressing for the control chamber was correlated with pressing for the stranger (females: R = 0.75, *p =* 0.013; males: R = 0.71, *p <* 0.005) but not with lever pressing for the partner.

The same cohorts were then tested for extinction of lever pressing over 10 days of trials in which the door was closed and the lever did not activate the motor. All groups extinguished lever pressing behavior within ~5 days of testing (Supplementary Figure S3). Repeated measures analysis revealed a significant effect of day of testing on pressing (F_(9)_ = 3.72, *p =* 0.0063) but no significant effect of the species or sex of the testing group (F_(2)_ = 0.76, *p =* 0.48).

## DISCUSSION

Male and female prairie voles worked for brief access to conspecifics, but exhibited qualitatively different patterns of pressing, indicating striking sex differences in social motivation. In females, lever pressing effort was based on familiarity of the social target (partner vs. stranger), but did not differ between same-sex (FF) and opposite-sex (FM) housed pairs. Because testing occurred with only the partner or stranger present at any given time, failure to spend time in the stranger chamber indicates lack of interest in the stranger, as opposed to relative preference for a better option. Females also exhibited extensive huddling with and time spent in the chamber of a partner but not a stranger. These preferences persisted when scaled by lever presses (i.e. time the subject was available), indicating strong selectivity in social preferences. Social motivation thus paralleled social selectivity in females.

Male prairie voles exhibited similarly strong selectivity in huddling and chamber preferences, consistent with decades of work showing partner preferences in both male and female prairie voles. In contrast, males showed no propensity to work harder to access a familiar vole than an unfamiliar social target, but instead worked significantly harder to access an opposite-sex social target than a same-sex social target. This reveals a dissociation between social motivation and markers of social bond formation such as huddling in males. This sex difference in motivated behavior is consistent with the hypothesis that outwardly similar partner preferences in males and females result from latent differences in underlying signaling pathways (De Vries, 2004). Oxytocin, dopamine, and opioid signaling all affect partner preferences in males and females (Williams et al., 1992a; Gingrich et al., 2000; Aragona et al., 2003; Burkett et al., 2011; Resendez et al., 2012; Johnson et al., 2016), but prairie voles also exhibit sex differences in these signaling pathways (e.g. Winslow et al., 1993; Martin et al., 2015; Ulloa et al., 2018). These latent differences in mechanisms underlying social bonding may support similar partner preference behavior while promoting sex differences in social motivation.

Factors that may particularly motivate males to access unfamiliar females include opportunities for mating and aggression. Male prairie voles exhibit multiple mating strategies in field settings, including both a socially monogamous “resident” partner strategy, and a “wanderer” strategy; however, even residents engage in extra-pair copulations (Madrid et al., 2020). Interest in non-partners can also result from motivation for aggressive behavior; for example, the opportunity for aggression is rewarding in dominant male mice (reviewed in Golden et al., 2019). That does not seem to be the case in male prairie voles, however. Aggression towards partners was rare, and response rate was not correlated to aggression towards a stranger in male prairie voles tested with females or males. Aggressive bouts in female prairie voles tested with female strangers initially appeared correlated with lever pressing effort, but this effect disappeared when scaled by access time, unlike effects reported for huddling/access time. When social interest is high (e.g. males for unfamiliar females), it is still possible that males would press more for their partners if placed in direct opposition to a stranger, and this is an avenue for future investigation.

### Oxytocin receptor signaling differs by relationship type and by individual social behaviors

Strong relationships were present between *Oxtr* genotype, OTR density, housing differences, and behavior, highlighting the role of genes and environment, and connections across levels of organiation. Oxytocin receptor density was strongly influenced by genotype in the nucleus accumbens and lateral septum, and was influenced by housing (opposite-sex vs. same-sex) in the other two brain regions sampled. Variation in OTR by NT204321 genotype replicated differences in the NAcc described in two prior reports (King et al., 2016; Ahern et al., 2021), and identified a relationship in the lateral septum that had not previously been reported. Variation in OTR density by relationship type has not been previously assessed, although oxytocin receptor density is known to be responsive to other housing manipulations such as early life family structure (Ahern and Young 2009), and solitary versus pair-housing (Pournajafi-Nazarloo et al., 2013, Prounis 2015), among other factors. Interestingly, genotype and housing were associated with neural receptor density changes in non-overlapping brain regions.

Oxytocin signaling plays a role in diverse social behaviors in prairie voles, including pair bond formation, consolation behavior, and alloparental care (Williams et al., 1992a; Olazábal and Young, 2006; Bales et al., 2007; Burkett et al., 2016). Furthermore, oxytocin signaling has been related to social reward in non-selective mice and hamsters (Dölen et al., 2013; Song et al., 2016; Borland et al., 2019). Strong correlations between nucleus accumbens OTR and lever pressing for the partner in the present study provide additional support for the role of nucleus accumbens OTR in social reward. Neural OTR was related to aggressive behavior as well as prosocial behavior, underscoring the complexity of oxytocin signaling in different brain regions (van Anders et al., 2013; Beery, 2015).

Genotype-behavior comparisons were examined in behaviors that could be pooled across housing conditions, which included aggressive behaviors in males and females, and lever pressing in females. Despite the relatively low presence of C carriers, there was a very strong correlation between *Oxtr* genotype and aggression in male prairie voles, again underscoring the connections between oxytocin and some “antisocial” social behaviors. In particular, while oxytocin promotes in-group social behaviors, enhanced selectivity for familiar individuals is accompanied by increased aggression towards strangers. This increased aggression is a well-described part of social bond formation in prairie voles (Bowler et al., 2002; Young et al., 2011; Resendez et al., 2012), and increased outgroup discrimination has also been described in humans (De Dreu et al., 2011).

### Species differences

Social pressing differed quantitatively but not qualitatively by species in meadow and prairie voles. Females of both species pressed more for partners than for strangers, but responses were lower in meadow voles, indicative of reduced social reward. This is consistent with prior findings from socially conditioned place preference tests, in which meadow voles did not condition towards a bedding associated with social contact, and in one setting conditioned away from it (Goodwin et al., 2019). These findings are also in line with results from the sole prior study of operant responses in voles. Matthews et al. (2013) tested prairie voles and meadow voles housed in long daylengths to determine whether they would learn to lever press for stranger voles. Only prairie voles demonstrated clear learning in this scenario, consistent with low stranger interest in meadow voles housed in the long day lengths that promote territorial behavior in this species (Beery et al., 2008b). Nonetheless, even under pro-social short daylength conditions, social pressing was low in meadow voles. Comparison of short daylength-housed female meadow vole responses for the partner chamber, stranger chamber, and an empty chamber in different trial blocks revealed equivalent levels of pressing for a partner or an empty chamber and less for the stranger. This suggests that decreased pressing for the stranger represents avoidance, but that pressing for the partner may indicate tolerance more than reward. Female (short daylength-housed) meadow voles also exhibited lower aggression than female prairie voles, consistent with social tolerance, and with prior descriptions of their behavior (Lee et al., 2019).

### Comparability across species and sexes

Lever pressing was demonstrated to be an effective metric to compare effort exerted to reach different social stimuli in voles; voles of each species and sex tested pressed at comparable rates for food reward, indicating a lack of major differences in task learning, and thus that social lever pressing can be assessed and compared across groups. Subject response rates were not consistently high or low across reward conditions, indicating that responses are reward-specific. Extinction was effective, with all subjects decreasing lever pressing behavior by more than half their baseline response count. Differences in lever pressing effort between groups could therefore be attributed to reward-specific differences in social motivation.

### Conclusions

While other studies have assessed social reward in rodents, few have considered the role of stimulus familiarity, likely because laboratory rodents do not exhibit familiarity preferences under normal conditions (reviewed in Beery and Shambaugh, 2021). In social choice tests, mice and young rats often prefer social novelty (Moy et al., 2004; Smith et al., 2015), and relative preference for a social stimulus versus a food stimulus is greater when novel rats are presented (Reppucci et al., 2020). Indeed, in operant trials in which rats had simultaneous access to familiar and unfamiliar same-sex conspecifics, rats expended more effort to access unfamiliar conspecifics (Hackenberg et al., 2021). In the present study, female prairie voles exhibited similar partner preferences but higher social motivation and aggression compared to female meadow voles. Social motivation and selectivity were not linked in male prairie voles, and there was a striking sex difference in the reward value of mates and peers in prairie voles. Oxytocin receptor genotype and receptor binding revealed connections between genotype, social environment, receptor density, and both prosocial and aggressive behaviors, showing the importance of this system across levels of biological organization. Better understanding of the interface between social motivation and social selectivity will thus be key to improving our understanding of the nature of social relationships.

## MATERIALS AND METHODS

### Animal subjects

Prairie voles and meadow voles from in-house colonies were bred in a long photoperiod (14h light:10h dark; lights off at 17:00 EST; described further in Lee et al., 2019). Meadow voles were weaned into the winter-like short photoperiods associated with group living in this species (10:14 light:dark; lights off at 17:00 EST). Voles were pair-housed in clear plastic cages with aspen bedding and an opaque plastic hiding tube. Food (5015 supplemented with rabbit chow; LabDiet, St. Louis, MO, USA) and water were provided *ad libitum*, except during food restriction (described below). All procedures adhered to federal and institutional guidelines and were approved by the Institutional Animal Care and Use Committee at Smith College.

### Timeline and groups

Training began in adulthood at 62±1.3 days of age (mean±SEM, range 41-76). Operant conditioning training and testing consisted of multiple phases described briefly here and in greater detail in subsequent sections. Responses (lever presses) were shaped and trained using a food reward on a fixed ratio 1 (FR-1) schedule. Animals that met training criteria progressed to the experimental testing sequence, beginning with 8 days of pressing for a food reward on a progressive ratio 1 (PR-1) schedule (Figure 1). Subjects in opposite-sex pairs were placed with an infertile but sexually active mate 5-10 days prior to the start of social habituation and testing. Subjects in same-sex pairs remained with their cage-mate. Social testing consisted of 8 days of PR-1 with rewards yielding access to the familiar (same- or opposite-sex) partner, and 8 days with access to a sex-matched stranger (order balanced within groups). A subset of voles continued in empty chamber control and/or extinguishing tests as described below. Voles were sacrificed at the conclusion of testing, and brains were stored at −80°C. Voles were trained and tested over 7 cohorts; group membership was distributed across cohorts.

We tested four groups of prairie voles (Figure 1): females lever pressing for a female conspecific (F➤F), females pressing for a male conspecific (F➤M), males pressing for a male conspecific (M➤M), and males pressing for a female conspecific (M➤F). Each group consisted of 8 focal voles, tested for 8 days with their partner and for 8 days with a series of novel “strangers”, sex-matched to the partner. The order of testing (partner then stranger or stranger then partner) was counterbalanced within groups. Some voles did not complete both partner and stranger testing, in which case additional voles were added up to 8/group. Meadow vole females (F➤F-*Mp*), n=7) were also trained and tested for 8 days of familiar and 8 days of novel vole exposure, with order counterbalanced within the group.

### Operant conditioning and testing with food reward

Subjects were weighed for three consecutive days to establish baseline body weights, then food-restricted to a target weight of 90% baseline to enhance motivation for the food reward. Weights were recorded daily after training or testing, prior to being returned to their home-cages. Any vole that dropped to or below 85% of the baseline weight was returned to *ad libitum* food to avoid long-term health consequences. Perforated cage dividers were used during food restriction to ensure each vole had access to its specific ration (0.3-1 food pellets and ~4g (half) of a baby carrot). Food restriction ended when subjects transitioned to social testing.

Operant conditioning was conducted in mouse-sized modular test chambers (30.5cm x 24.1cm x 21.0cm) outfitted with a response lever, clicker, modular pellet dispenser for mouse, and pellet receptacle (Med Associates Inc, St. Albans, VT, USA, Figure 1A). Data were acquired using the MED-PC-IV program running training protocols coded by experimenters. Sessions lasted 30 minutes and took place between 0900-1700. Vole behavior was shaped using manual reinforcement by an experimenter until a subject met the training criterion of three days in a row of ≥ five responses without manual reinforcement on an FR-1 schedule. One 20mg food pellet (Dustless Precision Pellet Rodent Grain Based Diet; Bio-Serv, Flemington, NJ, USA) was dispensed as each reward. Animals that did not learn to consistently lever press within ~20 days were used as partners or strangers for future social testing. Subjects that met the training criterion transitioned to a progressive ratio (PR-1) schedule with each successive reward requiring an additional response. The progressive ratio has been shown to be a better indicator of motivation than FR programs (Hodos and Kalman, 1963; Weatherly et al., 2003). PR-1 testing was conducted for 8 days, at the conclusion of which all focal animals were returned to *ad lib* food, and cage dividers were removed.

### Testing with social rewards

Social reward testing was conducted in mouse-sized modular test chambers, custom equipped with a motorized door (Med Associates Inc, St. Albans, VT, USA) for access to a second “social” chamber (Figure 1B). This chamber was constructed of clear plastic (15cm x 20.5 cm x 13cm) and contained an eye-bolt for tethering a stimulus vole (Figure 1C). A clear plastic tunnel (2.54 cm diameter, 5.5 cm long) connected the operant chamber to the social chamber, and the entire apparatus was fixed to a mounting board. Lever presses were rewarded by door opening and chamber access; the door remained raised for one minute, after which the experimenter returned the focal vole to the operant chamber. Sessions lasted 30 minutes and were video recorded for quantification of additional behaviors.

Subjects transitioned to social testing following a habituation session and two FR-1 sessions. Habituation to the social apparatus took place with the door open and the lever covered: voles explored the apparatus for 15 min with an empty social chamber, and 15 min with the partner tethered in the social chamber. Two days of FR-1 pressing for a tethered vole followed habituation to ensure that subjects associated lever pressing with access to the social chamber and a stimulus vole.

Social testing took part in two phases: pressing for a partner vole on a progressive ratio and pressing for a stranger on a progressive ratio. Each phase lasted 8 days. The order of testing was counterbalanced within groups and subjects completed both phases. Social stimulus animals were tethered to the end of the social chamber. During the eight days of stranger testing, the focal vole was tested against a novel vole each session to prevent familiarity between conspecifics.

### Non-social conditions

Empty chamber testing took place after social testing to avoid altering lever pressing for the social stimuli. The empty chamber control was run to assess the value of apparatus exploration: 30 voles (10 female prairie voles, 12 male prairie voles, 6 female meadow voles) pressed the lever for 8 successive days on a PR-1 schedule to access the adjacent chamber when no stimulus vole was present. Sessions lasted 30 minutes and video was recorded and scored for behavior after testing. For the extinction phase, 31 voles (13 female prairie voles, 11 male prairie voles, 7 female meadow voles) were tested in the social chamber with an unrewarded lever for 10 successive days (30 minute sessions).

### Behavioral scoring

Counts of responses (lever presses) and rewards (food pellets or door raises) were automatically recorded during each test. In all social trials (16/vole) and all empty chamber control trials (8/vole), behavior in the “social” chamber was also filmed with a portable digital video camera. Videos were scored using a custom perl script (OperantSocialTimer 1.1; A. Beery) to determine time in the social chamber, time in side-by-side contact with the tethered vole (huddling), and bouts of aggression. These values could also be reported relative to other intervals (e.g. time huddling/access time when the door was up, or time huddling/time in the social chamber). Non-social/empty chamber trials (8 days/vole) were also videotaped, and analyzed for time in the social chamber/available time with the door raised.

### Castration and tubal ligation

At least one week prior to pairing, the future “partner” of each opposite-sex prairie vole pair was surgically altered to prevent pregnancies during testing. Female partners of male focal voles underwent tubal ligation. Dorsal incisions were made over each ovary. Two knots were placed below each ovary at the top of the uterine horn. The wound was closed using a sterile suture. Male partners of female focal voles were castrated and implanted with testosterone capsules. Testes were accessed by midline incision, and the blood supply was cut-off through a tie at the testicular artery. Testes were removed and the muscle wall and skin were closed using sterile suture. A testosterone capsule was implanted subcutaneously between the scapulae. Capsules contained 4mm of crystalline testosterone (Sigma-Aldrich, St Louis, MO, USA) in silastic tubing (ID 1.98 mm, OD 3.18 mm; Dow Corning, Midland, MO, USA) as in (Costantini et al., 2007). Capsules were sealed with silicone, dried, and soaked in saline for 24h prior to insertion. A subset of strangers was also castrated or ligated, with no effect on focal behavior. Surgical procedures were performed under isoflurane anesthesia. Voles received 0.05mg/kg buprenorphine and 1.0mg/kg metacam subcutaneously prior to surgery, and again the following day. Post-operative wound checks continued for up to ten days post-surgery.

### *Oxtr* genotyping

DNA for NT213739 genotyping was isolated from frozen livers or cerebellar tissue using the Qiagen DNeasy Kit (Qiagen, #69506) using forward (5′ -CTCCTATTCAGCCCTCAGAAAC-3′) and reverse (5′ -TGAACCCTTGGTGAGGAAAC-3′) primers, exactly as described in (Ahern et al., 2021). These primers produce a 644 bp amplicon for which BsiHKAI cuts the C-allele to produce bands of 492 bp and 152 bp. Ilustra PuRe Taq Ready-to-Go PCR Beads (GE, #27-9557-01) were used with a PCR cycler (BioRad) set to 35 cycles (94° C denature, 55° C annealing, 72° C elongation) followed by a 1.5h BsiHKAI restriction digest prior to visualization using a 3% agarose gel (Hoefer, #GR140-500) infused with SYBR green and run for ~1h at ~100 V. DNA was analyzed from all prairie voles except the first cohort of six voles (n=30). Of these, 28 were successfully genotyped (14 females, 14 males). As in another prairie vole colony, the C/C genotype was rare (Ahern et al., 2021); thus C/C and C/T individuals were pooled as C carriers. Meadow vole genotype has not been previously assessed at this allele, and 8 females were tested as an initial sample. All were T/T, so no analysis of genotype-driven variation was warranted in that species.

### Receptor autoradiography

OTR binding density was assessed in the brains of 11 female prairie voles at the conclusion of the study (males were used for an additional pilot study). Frozen brains were sectioned coronally at 20μm, thaw-mounted on Super-frost Plus slides (Fisher, Inc.), and stored at −80° C until processing (as in Beery et al., 2008; Beery and Zucker, 2010; Mooney et al., 2015). Briefly, slides were thawed until dry, then fixed for 2 min in fresh, chilled 0.1% paraformaldehyde in 0.1 M PBS. Sections were rinsed 2×10 min in 50 mM Tris (pH 7.4), and incubated for 60 min at room temperature in a solution (50 mM Tris, 10 mM MgCl_2_, 0.1% BSA, 0.05% bacitracin, 50 pM radioligand) containing the radioactively labeled ^125^I-ornithine vasotocin analog vasotocin, d(CH_2_)_5_ [Tyr(Me)_2_,Thr^4^,Orn^8^,(^125^I)Tyr^9^-NH_2_] (^125^I-OVTA, PerkinElmer, Inc.). An adjacent series of slides, processed for non-specific binding, was incubated with an additional 50 nM non-radioactive ligand [Thr^4^Gly^7^]-oxytocin (Bachem). All slides were rinsed 3×5 min in chilled Tris– MgCl_2_ (50 mM Tris, 10 mM MgCl_2_, pH 7.4), dipped in cold distilled water, and air dried. Sections were apposed to Kodak BioMax MR film (Kodak, Rochester, NY, USA) for 3 days and subsequently developed. Radioligand binding density in each brain region was quantified in samples of uniform area from three adjacent sections for each brain region and averaged for each brain. Non-specific binding was subtracted from total binding to yield specific binding values.

### Statistical analyses

Social data were analyzed for all subjects completing both partner and stranger phases of testing (n=8 prairie vole M➤M pairs, 8 prairie vole M➤F pairs, 8 prairie vole F➤F pairs, 8 prairie voles F➤M pairs, and 7 meadow vole F➤F pairs). Four additional female prairie voles completed testing with a partner or stranger only: data from these subjects was included in analysis of food responses and food versus social response rates. Group differences in single variables (e.g. food responses) were assessed by 1-way ANOVA. 2-way repeated measures ANOVA (RM-ANOVA) was used to assess the effects of social factors, with stimulus familiarity [partner, stranger] as a within-subjects repeated measure, stimulus type [same-sex, opposite-sex] as a between-subjects (non-repeated) measure, a test for interaction effects [stimulus familiarity*stimulus type], and for subject matching. Paired *t*-tests were used within groups for comparison of behavior towards the partner vs. stranger. Response count (i.e. lever presses) and breakpoint (i.e. number of rewards achieved) are highly correlated; detailed results are therefore shown for only one measure (response count). Response rate (responses/active session time) was used when comparing food responses to social responses, as the lever was continuously active during food-rewarded testing (active session time = 30 min), but was not capable of raising the door when it was already up (active session time = 30-n minutes with the door up). Autoradiography data were collected in multiple brain regions, and comparisons were performed by 2-way ANOVA (group*brain region) followed by within-group comparisons adjusted using the False Discovery Rate procedure of Benjamini, Krieger, and Yekutieli (Benjamini et al., 2006). Statistical analyses were performed in JMP 15.0 (SAS, Inc.) and Prism 9 (GraphPad Software Inc.). All tests were two-tailed, and results were deemed significant at *p <* 0.05.

## AUTHOR CONTRIBUTIONS

AB, SL, and KB designed the experiments, SL, NB, and KB conducted the experiments, AB and NL performed receptor autoradiography, TA extracted DNA and genotyped the NT213739 SNP, AB and SL analyzed data and compiled the results, AB wrote the manuscript, and all authors edited the manuscript.

## ACKNOWLEDGEMENTS

We are extremely grateful to Dr. Tim Hackenberg for advising us on the design, physical setup, and analysis of the studies described here. Lab members Emily Halstead, Karina Lieb, Amelia Windorski, Rose Hatem, Madeleine Lerner and Marcela Rodrigues-Guimaraes assisted with behavioral testing and video scoring. Dale Renfrow (Smith Center for Design and Fabrication) assisted with building social testing chambers. We thank the staff of the Smith College Animal Care Facility for animal care and colony maintenance. This research was supported by the National Institute of Mental Health of the National Institutes of Health under Award Number R15MH113085.

---

**Figure S1.**
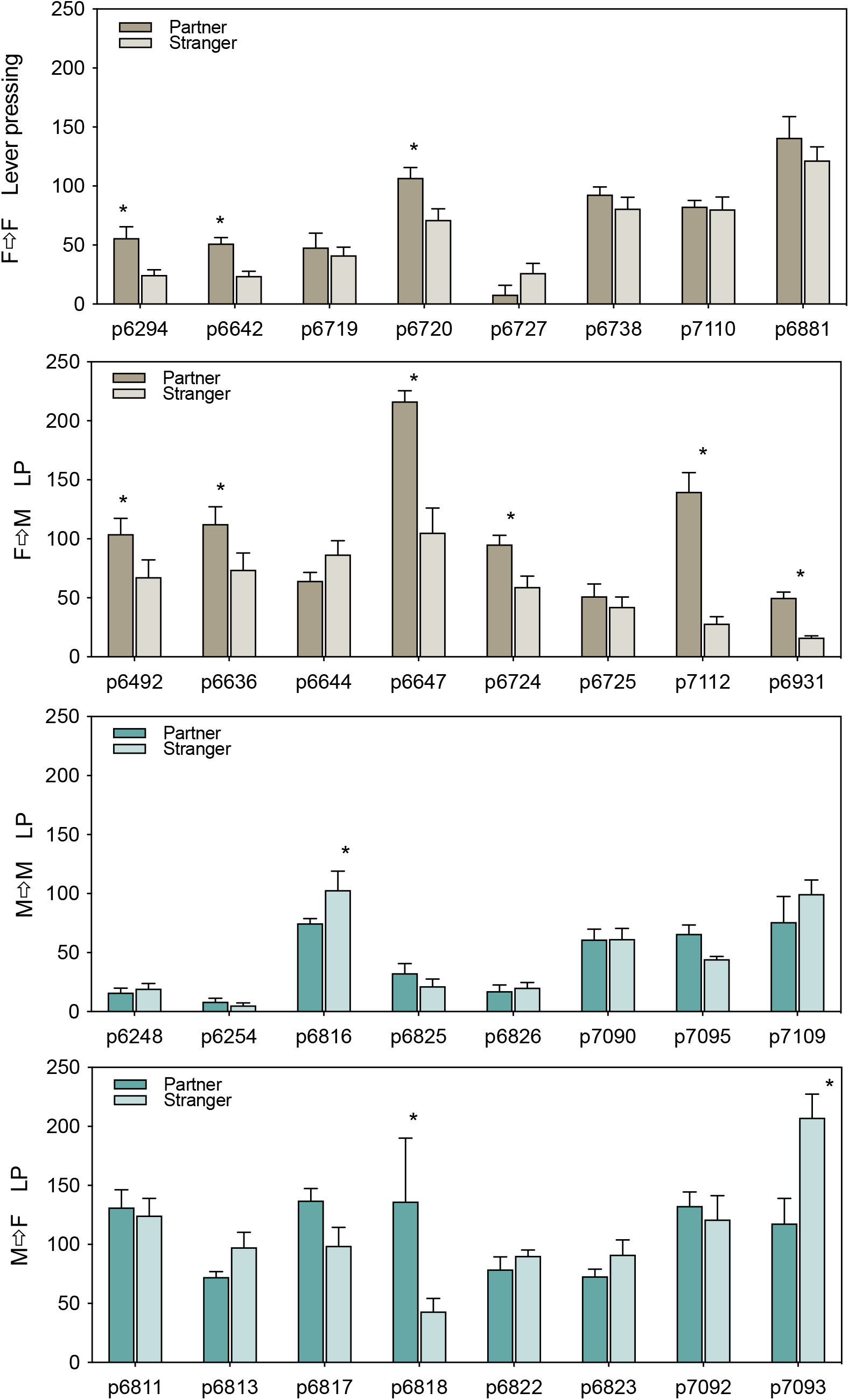
Individual lever pressing (LP) data for each prairie vole tested with a partner and stranger (8 days each). Significant within-individual preference for the partner>stranger was present more often in female

**Figure S2.**
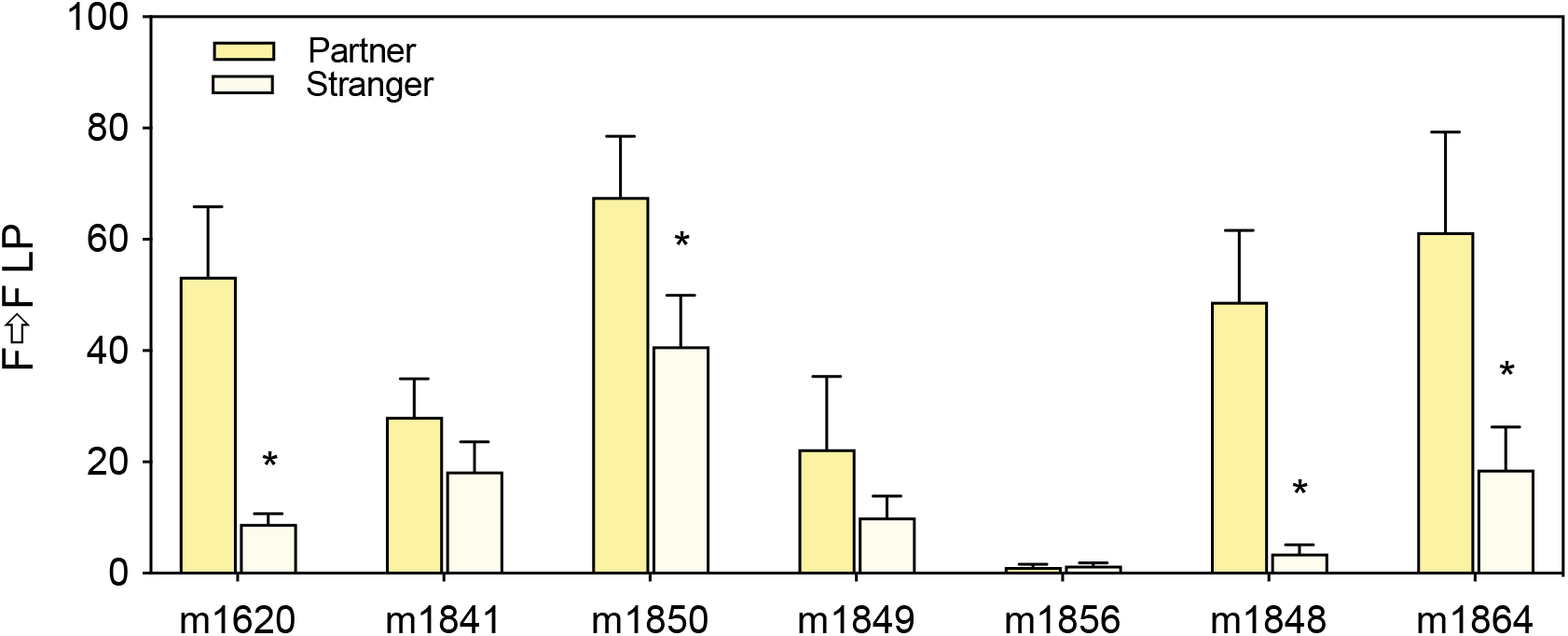
Individual data for each meadow vole tested with a partner and stranger (8 days each). Significant within-individual preference for the partner>stranger was detected in 4 of 7 females. One female did not press at high levels for any social stimulus, despite high levels of responding for a food reward during earlier testing.

**Figure S3.**
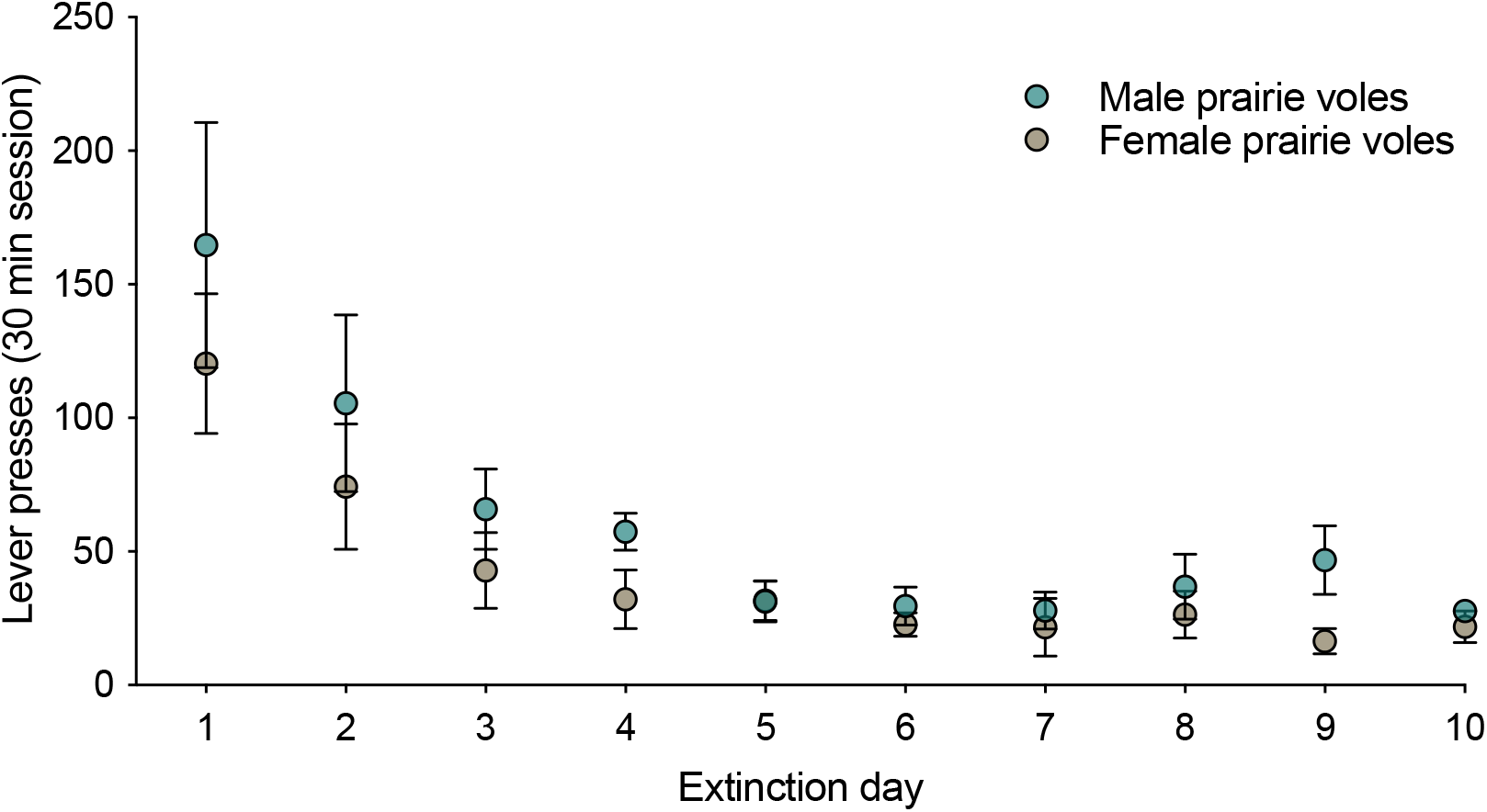
Extinction profile over ten days for each species and sex tested. Lever presses diminished rapidly over the first 4-5 days of testing with an inactive lever.

